# Instantaneous Phase-Shifting Optothermal Microscopy for Live-cell Metabolic Monitoring

**DOI:** 10.64898/2026.07.18.739313

**Authors:** Jiawang Qiu, Tao Yuan, Francesca Gasparin, Miguel A. Pleitez

## Abstract

Cellular metabolic activities can be studied using label-free vibrational spectroscopic imaging to leverage the endogenous contrast of biomolecules. However, fast live-cell imaging over large fields-of-view remains challenging due to the need for raster scanning and despite advances in wide-field modalities, imaging rapid cellular activities across large cell populations remains challenging. Here, we introduce a mid-infrared optothermal microscopy method termed Instantaneous Phase-Shifting Optothermal Microscopy (IPSOM). IPSOM circumvents conventional mechanical phase-shifting methods, achieved single-frame imaging speed 588-fold increase compared to the sequential approach, under field-of-view of 300×350 µm. For multi-wavelength hyperspectral imaging, IPSOM achieves an 8-fold speed improvement. IPSOM is used here to monitor lipid remodelling in adipocytes during lipolysis, demonstrating its potential for studying rapid cellular metabolism.

## Introduction

Vibrational spectroscopic microscopy (VSM) methods exploit the intrinsic differences in chemical-bond-selective absorption/scattering to achieve label-free molecular imaging. Hyperspectral VSM scanning generates image cubes combining spatial distribution with molecular absorption spectra, allowing researchers to distinguish between biomolecules. Representative examples include Coherent Anti-Stokes Raman Scattering (CARS) [1], Stimulated Raman Scattering (SRS) [2], and mid-infrared (mid-IR) absorption-based microscopy [3]. Most VSM methods rely on point-by-point scanning, which severely limits imaging speed and field of view (FOV). For instance, mid-IR optoacoustic microscopy (MiROM) [3] achieves high sensitivity with negligible photodamage but requires ∼1 minute to image a 50×50 µm FOV, and up to 7 minutes for a 2×2 mm area. To address the problem of slow image acquisition, computational frameworks like Bayesian raster-computed optoacoustic microscopy (BayROM) [4] has been introduced, generating images up to 10 times faster than conventional MiROM without prior training datasets. However, reconstruction-based methods for speed enhancement can introduce artifacts if the sample deviates from the assumed model. Consequently, point-scanning systems face a fundamental limitation: achieving a wide FOV requires prolonged acquisition times, while increasing imaging speed significantly reduces the FOV. This trade-off prevents the simultaneous monitoring of rapid biological events across large cell populations.

Several wide-field modalities have been developed to achieve imaging speed at large FOV’s. For instance, Bai et al. [5] introduced widefield photothermal sensing (WPS), achieving 1250 frames per second (fps) to map biomolecules in live cancer cells but at low signal-to-noise ratio (SNR). Additionally, Zhang et al. [6] developed bond-selective transient phase (BSTP) imaging, achieving an SNR of 100 at 1 fps with sub-micron resolution while other works integrated quantitative phase imaging (QPI) into optothermal modalities [7], or introduced synthetic-aperture variants [8] claiming spatial resolutions down to 120 nm in imaging of *E. coli* and *R. jostii*. However, while these wide-field modalities offer advantages such as high temporal resolution (WPS), improved SNR (BSTP), or high spatial resolution (QPI variants), they still struggle to simultaneously deliver rapid acquisition at large FOV’s. Recently, Yuan et al. [9] developed sequential phase-shifting optothermal microscopy (PSOM) to address the FOV limitation, achieving a 700×400 µm FOV with high phase sensitivity. This sensitivity enables the detection of weak optothermal signals from low-concentration molecules using low-power excitation, reducing phototoxicity. However, because this modality sequentially acquires multiple images at different phase-shift states, the mechanical switching mechanism limits the imaging speed to ∼0.017 fps [9].

To overcome the bottlenecks of sequential PSOM, we developed Instantaneous Phase-Shifting Optothermal Microscopy (IPSOM). Integrated with a polarization camera, IPSOM is capable of acquiring multiple phase-shifted images with a speed only limited by the exposure time of the camera. IPSOM achieves a raw imaging frame rate of 10 fps at a FOV of 300 × 350 μm, a 588-fold increase over state-of-the-art sequential PSOM (0.017 fps) [9]. For multi-wavelength hyperspectral imaging, IPSOM achieves 0.14 fps, an 8.2-fold improvement in acquisition speed [9]. Additionally, simultaneous acquisition intrinsically mitigates motion artifacts and environmental fluctuations, leading to a 3-fold improvement in contrast-to-noise ratio (CNR) while maintaining high structural similarity between the two imaging modalities (SSIM = 0.9761). Here, we demonstrate IPSOM’s ability to enable rapid label-free dynamic monitoring of lipid remodelling during lipolysis across large living cell populations.

## Results

### Optothermal Imaging Principle

As shown in Fig. 1A, IPSOM comprises two main units: a mid-IR excitation module and an instantaneous phase-shifting probe module. The excitation module generates a pump beam to stimulate molecular vibrations at the sample plane. It consists of an Optical Parametric Oscillator (OPO), a Germanium window to filter out visible light generated by the OPO, and a parabolic mirror that weakly focuses the mid-IR beam onto the sample area (300 × 350 µm).

**Figure 1.**
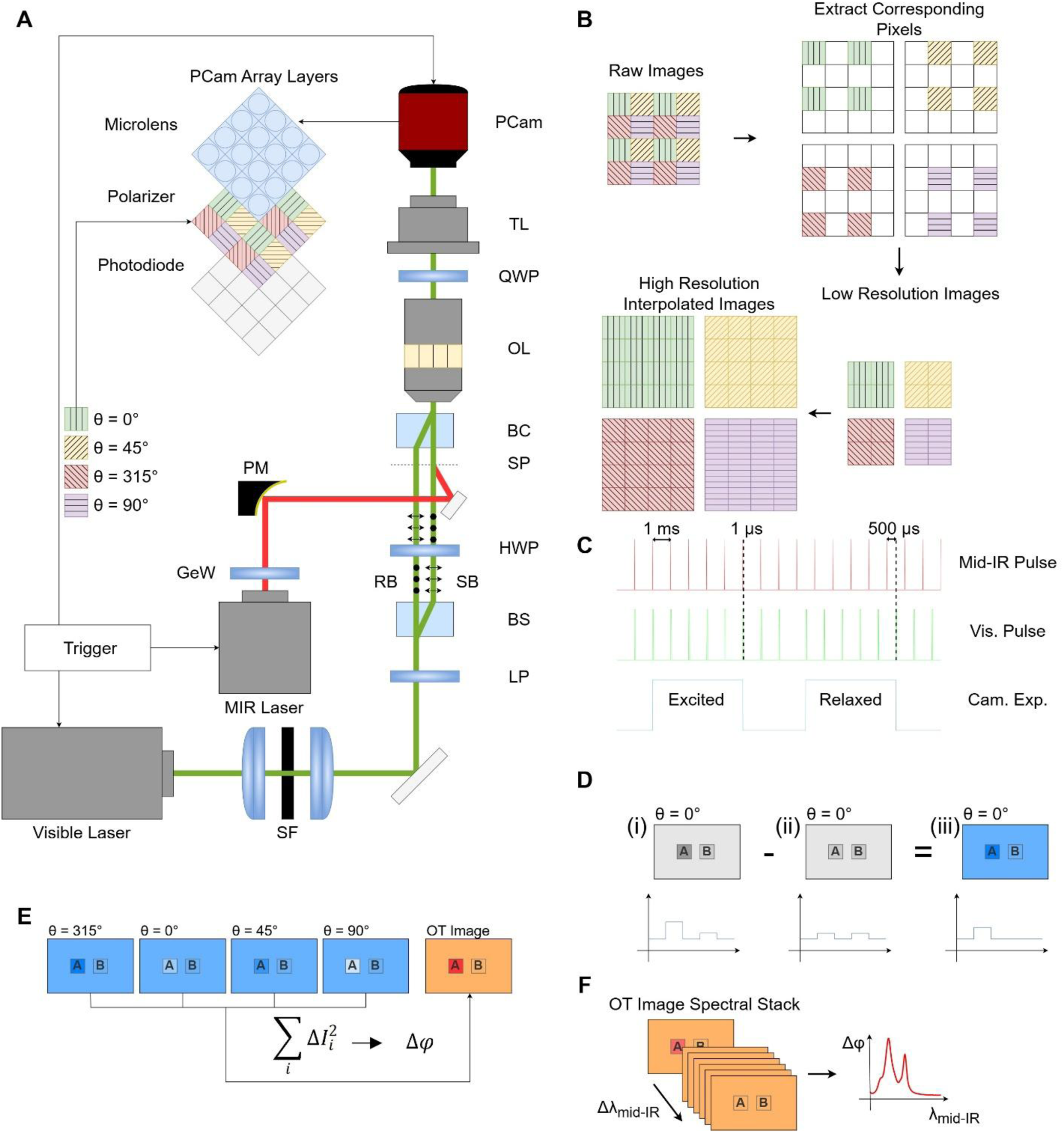
Imaging principle. **A.** System diagram. The system is built based on a phase-shifting quantitative phase imaging method and a pump and probe system to image the change in refractive index due to the mid-IR (pump) excitation pulse. A polarization camera is used to acquire the four phase shifting modules in parallel. SF: spatial filter, LP: linear polarizer, BS: birefringent beam splitter, HWP: half wave plate, SP: sample plane BC: birefringent beam combiner, OL: objective lens, QWP: quarter wave plate, TL: tube lens. GeW: Germanium window, PM: parabolic mirror, PCam: polarization camera, RB: reference beam, SB: sample beam. **B.** Preprocessing of polarization camera data. Interpolation is applied to improve the spatial resolution. **C.** Synchronization of the pump (mid-IR) and probe (visible) pulses and the camera exposure. **D.** Difference calculation between excited and non-excited states. (i) Excited state. (ii) Non-excited state. (iii) Subtraction image. **E.** The square sum of the images obtained at the four different polarization angles are used to quantitatively calculate the change in phase due to the mid-IR excitation. **F.** Stack of images are collected along a selected wavelength range to obtain mid-IR induced OT chemical spectrum.

The probe module employs an interferometric system where the input beam is split into two orthogonal polarization beams: a sample beam (SB) and a reference beam (RB). These beams are recombined after traversing the sample plane to form a balanced Mach-Zehnder interferometer. The phase difference between the SB and RB induces changes in optical intensity detected by the camera sensor array. Detailed descriptions of phase calculation using Phase-Shifting Microscopy (PSM) at different analyser polarization angles can be found in Yuan et al. [9]. However, unlike previously reported, [9] IPSOM does not require mechanical rotation of the analyser to sequentially obtain PSM images, which significantly increases imaging speed. Instead, in IPSOM, as illustrated in Fig. 1A and B, a polarization grid featuring four distinct polarizer orientations is mounted directly over the camera sensor. This allows for the simultaneous capture of four PSM images at different polarization angles within a single frame. This sequential acquisition configuration reduces low frequency noise, including fluctuations in laser energy, optical path changes due to mechanical movement, and environmental disturbances. Bicubic interpolation is subsequently applied to reconstruct missing pixels between the polarized regions.

Due to the rapid relaxation of the optothermal signal in water (32.8 µs) [10], precise synchronization between camera exposure, mid-IR pump pulses, and visible probe pulses is critical. As shown in Fig. 1C, the synchronization is divided into two stages: excited and relaxed. During **excited acquisition**, the visible pulse is generated approximately 1 µs after the mid-IR pulse to capture the refractive index change induced by molecular excitation. During **relaxed acquisition**, the probe pulse arrives 500 µs after the mid-IR pulse, capturing an image after the molecular relaxation process is complete (i.e., a baseline with negligible phase change). The mid-IR-induced phase change is calculated by subtracting the excited image from the relaxed image (Fig. 1D). These subtraction images at four different polarization angles are combined using **Equation (1)** (see Methods) to generate the final IPSOM image (Fig. 1E).

### Validation of the Instantaneous Phase-Shifting imaging

To validate the instantaneous phase-shifting method and demonstrate its potential imaging speed, we compared its quantitative phase imaging (QPI) capability with conventional sequential phase-shifting method. The comparison results are presented in Fig. 2. First, we evaluated quantitative phase retrieval using a focus star phase target (nominal height: 150 nm). Fig. 2A and B show the phase-shifting microscopy (PSM) images acquired by instantaneous method and sequential method for polarizer angles of (i) 315° and (ii) 0°, respectively; PSM images for the remaining angles (45° and 90°) are provided in Supplementary Fig. 1. Fig. 2C and D display the resulting quantitative phase images (QPIs), calculated using phase-shifting algorithm, as shown in **Equation (2)** (see methods).

**Figure 2.**
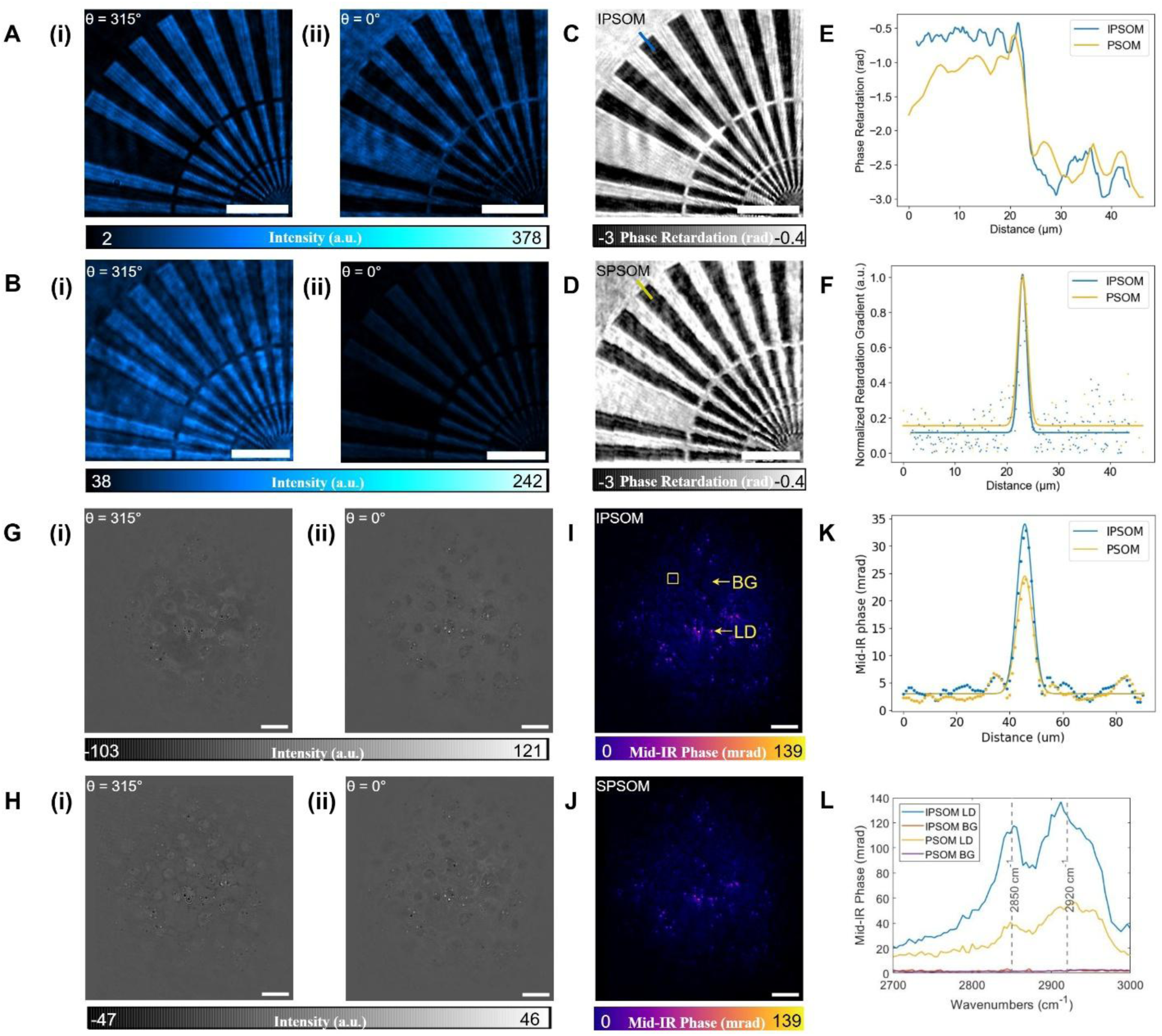
Validation of Instantaneous Phase-Shifting Optothermal Microscopy (IPSOM). IPSOM images of a focus star (feature height 150 nm) and fixed adipocytes (day 6 differentiation) are compared to the PSOM method to prove that IPSOM can acquire comparable mid-IR phase images with conventional methods. **A.** PSM images of a focus star acquired via IPSOM at polarization angles of 315° (i) and 0° (ii). **B.** PSM images of a focus star acquired via PSOM at polarization angles of 315° (i) and 0° (ii). **C.** Phase retardation image of a focus star acquired using IPSOM. **D.** Phase retardation image acquired using PSOM. **E.** Line profile over the edge of the focus star, highlighted by blue and yellow lines in **C** and **D**. **F.** Gaussian fit of the LSF obtained from **E**, the resolution, 1.83 µm for IPSOM and 2.18 µm for PSOM, is estimated using the FWHM of the fitted LSF. **G.** PSM images of differentiated adipocytes acquired via IPSOM at polarization angles of 315° (i) and 0° (ii). **H.** PSM images of differentiated adipocytes acquired via PSOM at polarization angles of 315° (i) and 0° (ii). **I.** Corresponding mid-IR phase image at 2850 cm^-1^ acquired using IPSOM. **J.** Corresponding mid-IR phase image at 2850 cm^-1^ acquired using PSOM. **K.** Line profile over a lipid droplet highlighted by the yellow square in **I**. **L.** Spectra at the highlighted lipid droplet (LD) and background (BG) in **I**, comparison between the spectra of IPSOM and PSOM shows a 3-fold increase of the mid-IR phase signal at the CH_2_ absorption peaks (2850 cm^-1^ and 2920 cm^-1^). Scale bars: 50 µm.

The instantaneous-QPI (I-QPI) image exhibits a finer projected pixel size (0.21 µm) compared to the sequential-QPI (S-QPI) image (0.7 µm/pixel), though the polarization grid initially reduces effective spatial sampling density; bicubic interpolation restores the full field. Fig. 2E shows line profiles across the edge of the focus star (corresponding to the blue and yellow lines in Fig. 2C and D). The instantaneous method yields results that agrees with the sequential results, despite stronger fluctuations in S-QPI. We derived the line spread function (LSF) by taking the first derivatives of the line profiles in Fig. 2E; the Gaussian-fitted LSF is shown in Fig. 2F. The resolution was evaluated using the full width at half maximum (FWHM) of the fitted LSF, yielding 1.83 µm for I-QPI and 2.18 µm for S-QPI. I-QPI demonstrates higher resolution attributed to improved spatial sampling rather than a change in the diffraction limit.

We then incorporated the excitation module to verify IPSOM’s capability for chemical bond-specific imaging. For this purpose, we compared mid-IR phase imaging results from both modalities (sequential and instantaneous method) using fixed (day-6 differentiated) adipocytes imaged at 2850 cm^-1^. Fig. 2G and H show the PSM subtraction images at polarizer angles of (i) 315° and (ii) 0° (details in Supplementary Fig. 1), while the resulting quantitative mid-IR phase images are shown in Fig. 2I and J. We used the Structural Similarity Index Measure (SSIM) to quantify similarity between the PSM subtraction images of both modalities. The SSIM values for the PSM subtraction images were 0.9722, 0.9728, 0.9778, and 0.9642 for polarizer angles of 315°, 0°, 45°, and 90°, respectively; the SSIM for the final mid-IR phase image was 0.9761. Fig. 2K shows a line profile across the lipid droplet highlighted by the yellow square in Fig. 2I, zoomed-in versions are shown in Supplementary Fig. 2. The similar line profiles further confirm high concordance between the two methods. Fig. 2L presents spectra acquired from the lipid droplet (LD) and background region (BG), highlighted by arrows in Fig. 2I (see zoomed-in of Supplementary Fig. 2). The LD signal at CH₂ absorption peaks (2850 cm^-1^ and 2920 cm^-1^) increased by approximately three-fold for IPSOM compared to PSOM, while the BG signal remained the same. This resulted in a 3.2-fold improvement in Contrast-to-Noise Ratio (CNR) while using the same excitation power. Most importantly, the imaging frame rate was 10 fps for IPSOM compared to 0.012 fps for PSOM, representing an 833-fold improvement in imaging speed.

### Chemical Specificity and Quantitative System Validation

To demonstrate that chemical contrast images can be obtained using IPSOM, we acquired a hyperspectral image stack of fixed differentiated adipocytes (day 6) containing mature lipid droplets. The results are presented in Fig. 3A–F. Fig. 3A displays the imaged region captured via a Zernike Phase Contrast (ZPC) microscope. Fig. 3B – E present IPSOM images at wavenumbers of 2850 cm-1, 2920 cm-1, and 2700 cm-1 along with an image acquired without mid-IR excitation. As show in Fig. 3B and Fig. 3C, the CH_2_ symmetric and asymmetric stretching vibrations within lipid droplets exhibit signal on 2850 cm^-1^ and 2920 cm^-1^. Residual signal in the 2700 cm^-1^ image (Fig. 3D) might arise from non-specific background absorption (such as water and the overall cytoplasm) rather than lipid-specific contrast. Nevertheless, signal strength and contrast for the lipid droplets are significantly higher at 2850 cm^-1^ and 2920 cm^-1^ compared to 2700 cm^-1^, confirming that IPSOM can distinguish lipid molecules. To further validate, we extracted IPSOM spectra from both the lipid droplets and the background medium (Fig. 3F); a normalized FTIR spectrum of triglyceride (TAG) is included as a reference. The IPSOM spectrum of the lipids exhibits peaks consistent with the FTIR reference TAG spectrum.

**Figure 3.**
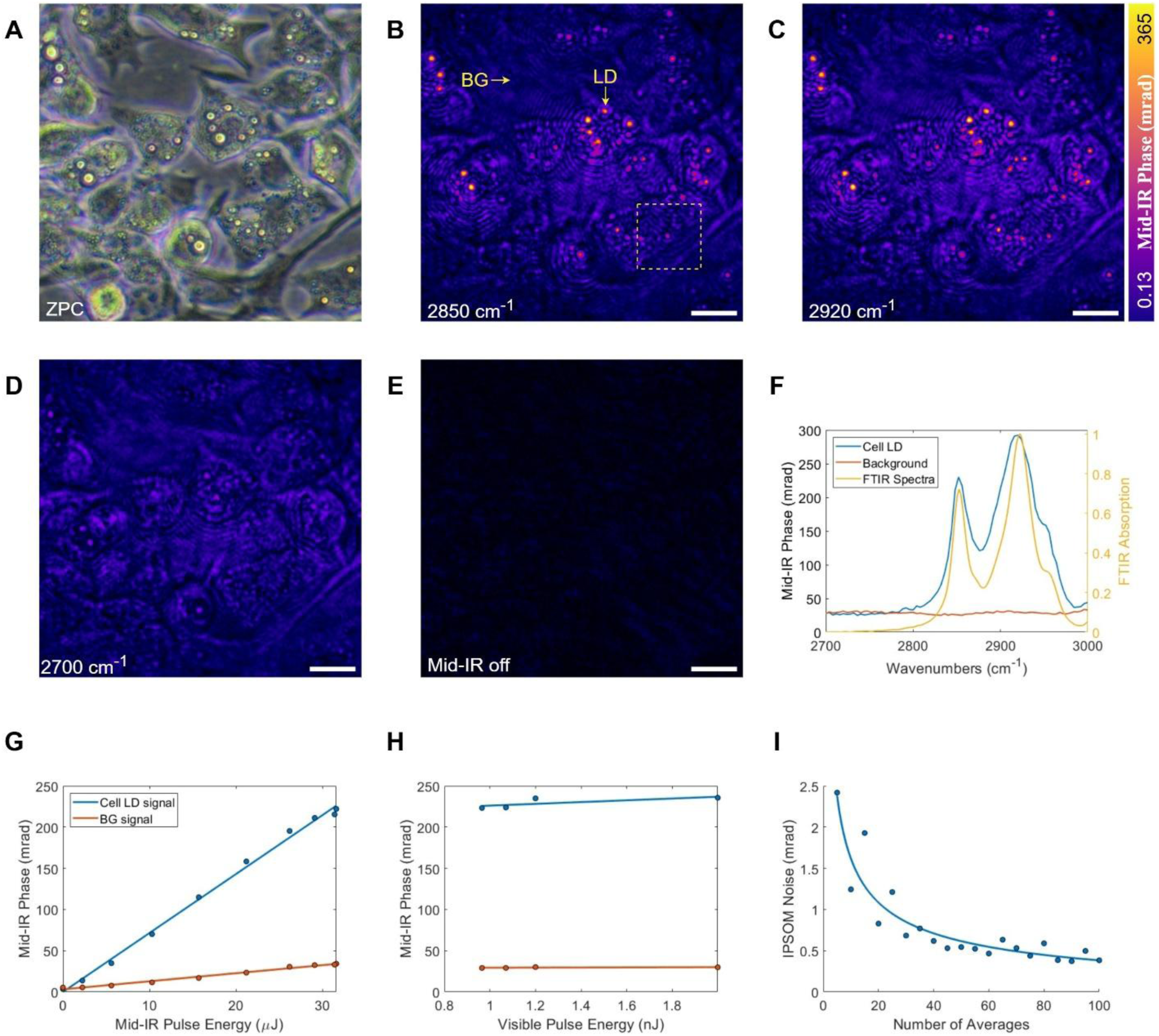
Hyperspectral imaging proof of concept. **A.** Zernike Phase Contrast (ZPC) microscope image of fixed differentiated (day 6) adipocytes. **B-E.** Instantaneous Phase-Shifting Optothermal Microscopy (IPSOM) mid-IR phase images of the region shown in **A**, at wavenumbers 2850 cm^-1^ (**B**), 2920 cm^-1^ (**C**), 2780 cm^-1^ (**D**) and with mid-IR beam blocked (**E**). **F.** IPSOM spectra of background (BG) and lipid droplet inside adipocyte (LD) as indicated in **B**, yellow line shows comparison with the FTIR spectra of TAG used as reference. **G.** IPSOM signal dependency to mid-IR pulse energy at 2850 cm^-1^. **H.** IPSOM signal dependency to 532 nm visible pulse energy, with mid-IR laser set to 2850 cm^-1^. **I.** Correlation between IPSOM noise power and number of images used for averaging. Scale bars: 25 µm.

To systematically characterize the quantitative performance and operational robustness of IPSOM, we conducted three complementary measurements (Fig. 3G–I). Fig. 3G evaluates the dependence of the measured mid-IR phase on the excited pulse energy. The response exhibits a strict linear relationship across the operational range (linear fitting slope: 7.16 mrad/µJ, R^2^: 0.996), confirming that IPSOM faithfully preserves the linear absorption-to-phase conversion dictated by optothermal theory [6]. The linearity of IPSOM is a critical advantage over nonlinear scattering-based modalities [8], as it gives the potential for direct, quantitative mapping of local chemical concentrations without empirical corrections. In Fig. 3H, we assessed the stability of mid-IR phase with respect to probe pulse energy. It is observed that the detected mid-IR phase for both LD and background remain invariant with respect to probe energy varying from 0.97 to 2.0 nJ, demonstrating that the IPSOM measurement is decoupled from probe intensity instability. This inherent robustness eliminates the need for active stabilization or external feedback loops, simplifying system alignment and enhancing long-term experimental stability. In Fig. 3I, we characterized the noise performance as a function of temporal averaging. The root-mean-square noise scales as *N*^−0.515^, adhering to theoretical shot-noise-limited statistics *N*^−1/2^. The result also suggests that averaging ∼30 frames can provide a moderate enhancement in contrast-to-noise ratio.

### Longitudinal Live-Cell Functional Monitoring

A key advantage of IPSOM is its capacity for label-free hyperspectral imaging of living cells with exceptional temporal stability. Unlike fixed-cell assays that capture static snapshots, longitudinal metabolic tracking requires uninterrupted, signal-drift-free acquisition to resolve continuous biochemical transitions. To evaluate IPSOM’s suitability for extended live-cell monitoring, we acquired hyperspectral image cubes over a 200 min period at a 9-min temporal resolution. Representative fields of view at the CH₂ stretching resonance (2850 cm^-1^) and an off-resonance reference (2700 cm^-1^) are shown in Fig. 4A and B, with the corresponding molecular spectra from lipid droplets (LDs) and the surrounding cytoplasm plotted in Fig. 4C. Across the extended acquisition, spectral traces at identical spatial locations exhibited remarkable stability, with phase fluctuations constrained to ∼2 mrad (average peak-to-peak) for both LDs and background regions (Fig. 4D and E). This long-term stability is intrinsically linked to IPSOM’s instantaneous phase-shifting architecture, which eliminates mechanical hysteresis and suppresses low-frequency environmental drift. Such temporal fidelity is critical for resolving slow but physiologically significant metabolic processes, including lipogenesis and lipolysis, which typically unfold over multi-hour timescales.

**Fig. 4.**
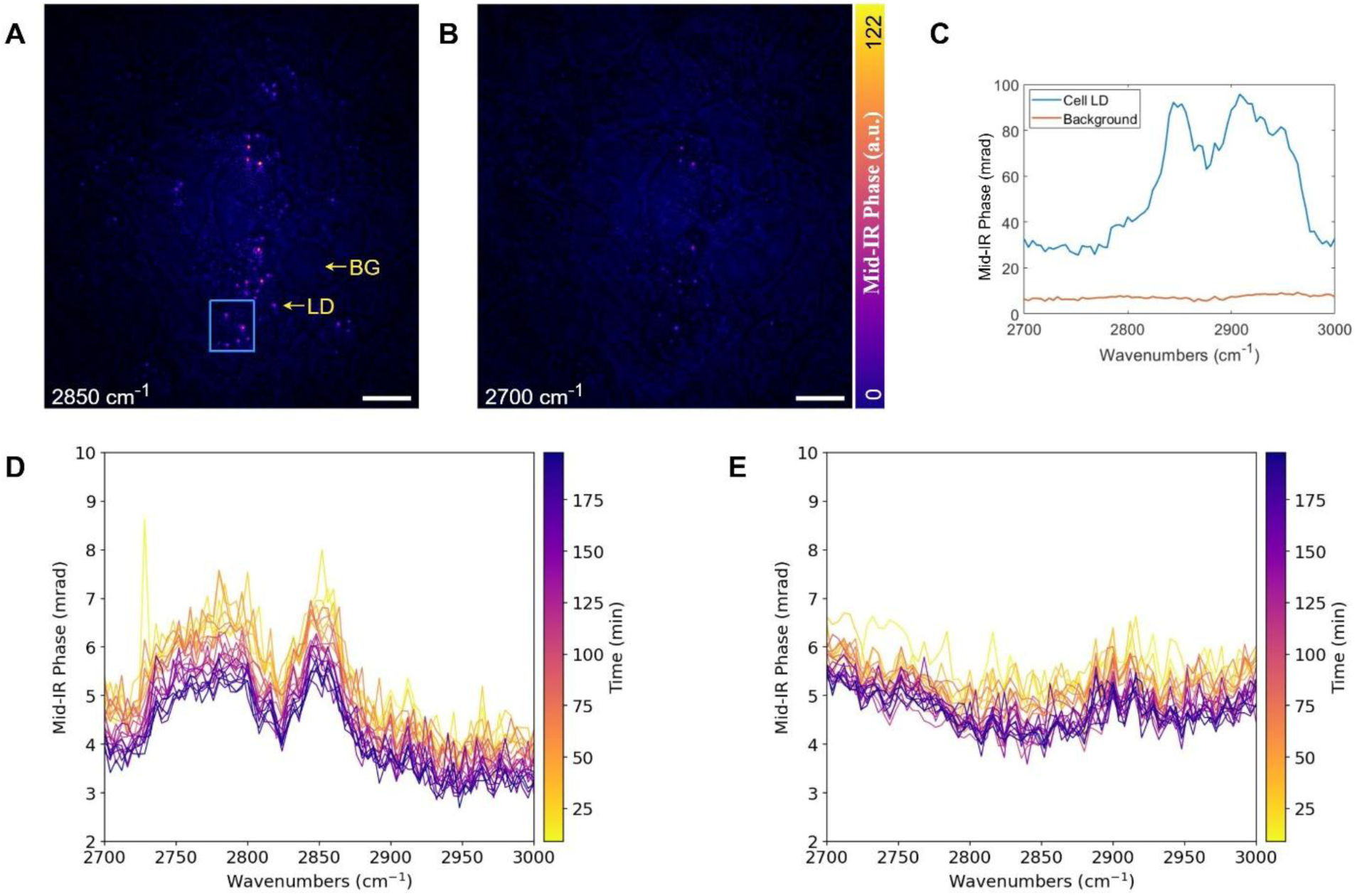
Hyperspectral live-cell imaging results. Continuous hyperspectral images of live differentiated (day 6) adipocytes are acquired. **A.** Image acquired at 2850 cm^-1^. **B.** Image acquired at 2700 cm^-1^. **C.** Comparison between the OT spectra of lipid droplets inside cells and background, as indicated by LD and BG arrows in **A**. **D.** Change in spectra over time at the squared region in **A**. **E.** Change in spectra over time of the background area. Scale bars: 50 µm

Building on this stability, we next demonstrated IPSOM’s capacity to non-invasively track dynamic lipid remodelling in live adipocytes. The synthesis, storage, and mobilization of TAG are central to cellular energy homeostasis, yet their real-time quantification in living populations remains challenging without perturbing labels [11]. To address this, we stimulated lipolysis in day-6 differentiated 3T3-L1 adipocytes with 10 µM forskolin and monitored lipid droplet dynamics at 2850 cm^-1^ over 400 min. Time-series IPSOM images (Fig. 5A-G) reveal progressive LD shrinkage and eventual resorption following drug administration. Quantification of LD signal (Fig. 5H) showed a sustained reduction in lipid signal to ∼35% of baseline, in stark contrast to control experiments, which exhibited minimal lipid redistribution (Supplementary Fig. 3). Together, these results establish IPSOM as a robust, label-free platform for high-throughput, longitudinal metabolic phenotyping, bridging the gap between rapid optothermal imaging and physiological relevance.

**Figure 5.**
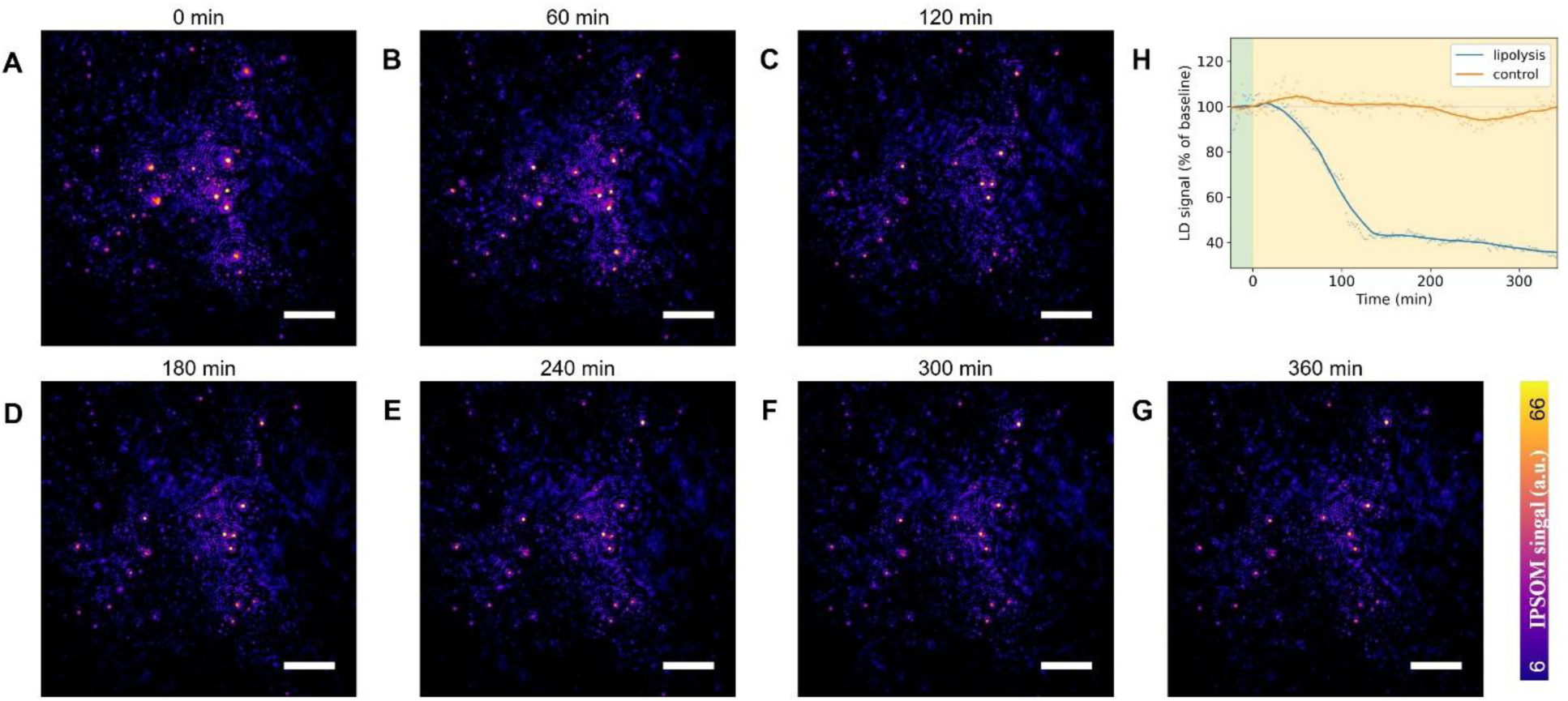
Instantaneous Phase-Shifting Optothermal Microscopy (IPSOM) live-cell imaging – monitoring lipid remodelling during lipolysis and change in membrane potential during membrane depolarization. The adipocytes are stimulated with 10 µM forskolin to induce lipolysis and monitored over ∼400 minutes. **A-G.** Representative time series of IPSOM images of live differentiated (day 6) adipocytes acquired at 2850 cm^-1^. **H.** Lipid droplet (LD) signal, comparison between control and stimulated, decrease of ∼65% is observed during lipolysis. Scale bars: 25 µm.

## Discussion

In this work, we introduced IPSOM, a label-free live-cell imaging platform that simultaneously delivers molecular specificity, fast acquisition speed and large FOV. Unlike fluorescent lipid probes such as Nile Red [12] and autofluorescence-based fluorescence-lifetime imaging microscopy (FLIM), which either require exogenous staining or yield only indirect metabolic readouts without molecular specificity, IPSOM exploits intrinsic molecular vibrational contrast for acquiring chemical-specific images without perturbing cell physiology, achieving molecular-specific imaging at 10 fps across large living cell populations at FOVs up to 300 × 350 µm.

Specifically, by simultaneously acquiring PSM images at different polarization angles, IPSOM achieved a raw imaging frame rate of 10 fps – a 588-fold increase over state-of-the-art sequential PSOM (0.017 fps) [9]. For continuous measurements at a single wavelength, the speed is 41.7 times faster than traditional sequential PSOM. For multi-wavelength hyperspectral imaging, the obtained imaging speed is 0.14 fps representing an 8.2 times improvement [9]. Additionally, the simultaneous acquisition mechanism intrinsically mitigates motion artifacts and environmental fluctuations, yielding a 3-fold enhancement in the contrast-to-noise ratio (CNR), while maintaining high similarity (SSIM = 0.9761) compared to the established sequential PSOM. This methodological advancement overcomes the compromise between imaging speed and FOV in VSM, enabling the capture of transient biological events that were previously impossible to monitor label-free in wide-field.

IPSOM was successfully applied for continuous, label-free monitoring of rapid metabolic activities in living cells, specifically, we tracked lipid remodelling during lipolysis in differentiated 3T3-L1 adipocytes. Unlike conventional methods that rely on phototoxic fluorescent markers to observe lipolysis and lipid remodelling, IPSOM provides a non-destructive alternative that enables the observation of native metabolic flux. This allows for the observation of native biological responses to drug interference across larger cell populations, providing deeper insights into cell metabolic properties.

Overall, IPSOM establishes a high-speed, large-FOV label-free imaging platform, enabling the study of fast metabolic activities in large cell groups. Further developments, in particular increasing camera acquisition speed (currently at 20 fps) will boost for hyperspectral imaging speed. Additionally, the use of a high numerical aperture (NA) objectives (currently NA of 0.28) will enhance spatial resolution and achieve refraction-limit resolution of the used probe beam (i.e. ∼250 nm). Furthermore, the reported spectral range is limited to the C-H stretching region (2800–3000 cm-1) and could be extended towards the fingerprint region to include protein and carbohydrate signatures.

In summary, we have demonstrated that IPSOM overcomes the traditional trade-offs between imaging speed and FOV in VSM by leveraging single-shot optothermal imaging. This approach offers a powerful, non-invasive tool for biomedical research, and we anticipate its broad application in studying fast metabolic interactions within large cell populations.

## Methods

### IPSOM setup

The IPSOM system was built using a synchronized pump-probe architecture. For the probe channel, a 532-nm pulsed laser (Cobolt 06-Tor, HÜBNER GmbH & Co. KG) was operated at a 2-kHz repetition rate with an initial linear polarization of 90° relative to the x-axis. The pump channel utilized an optical parametric oscillator (OPO NT277, EKSPLA) pulsing at 1 kHz. To isolate the mid-IR signal, a germanium window was placed after the OPO output to filter visible light originated from the OPO, and a rotating polarizer in the mid-IR range was included for manual adjustment of the pump power. This beam was weakly focused onto the sample plane by a parabolic mirror with a numerical aperture (NA) of 0.062 (Edmund Optics Ltd., #37248).

In the probe beam path, a 45° polarized beam was first generated using a linear polarizer (Thorlabs Inc., LPVISC050MP2). This light was partitioned into two orthogonal components— a 0° sample beam (SB) and a 90° reference beam (RB)—by a 1-mm polarizing beam displacer (Newlight Photonics Inc., PDB1010-AR800/400). A half-wave plate (Thorlabs Inc., WPHSM05-532), oriented with its fast axis at 45°, shifted these polarizations by 90°, resulting in an SB at 90° and an RB at 0°. While the SB was directed through the sample, the RB passed through an adjacent clear area.

After recombination via a polarizing beam combiner, the light was collected using a 10× objective (NA = 0.28, Thorlabs Inc., MY10X803). The signal was then modulated by a quarter-wave plate (Thorlabs Inc., WPQSM05-532, fast axis at 45°) and focused by a tube lens (MXA22018) onto a polarization-sensitive camera (Thorlabs Inc., CS505MUP). Timing for both the pump and probe triggers was managed and synchronized through a Texas Instruments MSP430F5529 LaunchPad microcontroller.

### 3T3-L1 Adipocyte Culture and Differentiation

Mouse 3T3-L1 fibroblasts (ATCC CRL-1658) were cultured in custom imaging dishes and maintained in standard growth medium: low-glucose DMEM (1 g/L glucose; Merck) containing 10% fetal bovine serum (FBS; Merck) and 1% penicillin-streptomycin (Merck). Upon reaching full confluence, adipogenic differentiation was initiated using medium consisted of high-glucose DMEM (4.5 g/L glucose; Merck), 10% FBS, 1% penicillin-streptomycin, 1 μg/mL insulin (Sigma-Aldrich), 0.25 μM dexamethasone (Sigma-Aldrich), 0.5 mM 3-isobutyl-1-methylxanthine (IBMX; Sigma-Aldrich), and ABP (1:1000 dilution of 50 mg/mL l-ascorbate, 1 mM biotin, and 17 mM pantothenate; Sigma-Aldrich). The full differentiation was applied on differentiation days 0 and 2. On day 4, cells received differentiation medium containing only insulin (1 μg/mL) and the ABP supplements. On day 6, cells were returned to standard growth medium (low-glucose DMEM). Throughout differentiation and imaging, cells were maintained at 37°C in a humidified 5% CO2 incubator. All IPSOM measurements were performed in growth medium.

### TAG phantom preparation

For calibration experiments, synthetic triglyceride (TAG) droplets were prepared by dissolving 1 mg of 1,2-dioleoyl-3-palmitoyl-rac-glycerol (Sigma-Aldrich) in 100 μL of 2:1 chloroform:methanol to yield a 10 mg/mL suspension. A 10 μL aliquot of this suspension was deposited onto the ZnS imaging window of a custom dish and allowed to air-dry at room temperature until complete solvent evaporation. The dish was then filled with deionized water to a depth of 3 mm and sealed with a cover glass.

### Monitoring lipolysis

Day 6 differentiated 3T3-L1 adipocytes were imaged at room temperature using an IPSOM system with a pump pulse energy of 15.7 µJ. After recording the baseline for 25 minutes, 10 µM forskolin (Sigma-Aldrich), obtained by dissolving forskolin powder in dimethyl sulfoxide, was added to induce lipolysis. The cells were monitored for a total duration of 400 minutes.

### Image processing

Pixels corresponding to polarization angles of 315°, 0°, 45°, and 90° were extracted, and interpolation was applied to improve resolution.

For mid-IR phase imaging, non-excited images were subtracted from excited images. Background subtraction was applied to enhance visual contrast. The signals were combined using:

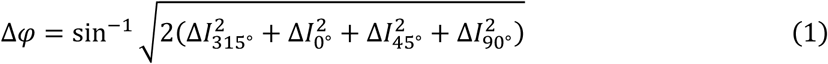

where *Δ*φ is the mid-IR induced local phase perturbation (or mid-IR phase), 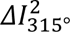 is the subtraction image from the 315° polarizer grids.

For acquiring quantitative phase retardation (QPI), PSM data from the four polarization angles were combined using:

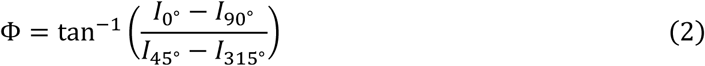

where Φ is the phase retardation.

Finally, PSF deconvolution was performed on the mid-IR phase data to compensate for the Airy disk and other artifacts, details about the PSF deconvolution results can be found in Supplementary Fig. 4. Lipolysis quantification used IPSOM images prior to PSF deconvolution.

### Lipid Signal Quantification

IPSOM image stacks were corrected for slow instrumental drift by fitting a third-degree polynomial to the per-frame mean of the paired non-excited images and dividing each frame by the trend.

LD signal was quantified per frame using a threshold derived from the pixel intensity histogram. The background distribution was modelled as a Gaussian fit to the peak-side dense region, and an intensity threshold separating background from LD signal was obtained from this fit. The LD signal metric was the sum of IPSOM intensities over all pixels exceeding this threshold, expressed as percentage change from the pre-stimulation reference frame after rolling-mean smoothing.

### Evaluation and Validation

Image quality was quantitatively assessed using the SSIM, noise analysis, SNR and CNR.

The Structural Similarity Index (SSIM) was calculated to measure the similarity between the IPSOM image x and the reference PSOM image y, based on luminance, contrast, and structure:

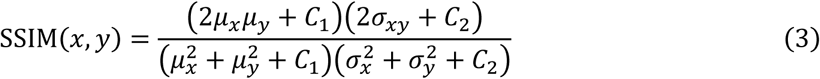

Where μx and μy are the pixel sample means, 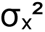 and 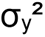 are the variances, σ_xy_ is the covariance, and C1 and C2 are constants to stabilize division with weak denominators.

Noise was quantified as the standard deviation (σ) of pixel intensities of the image acquired when the mid-IR pump beam is blocked:

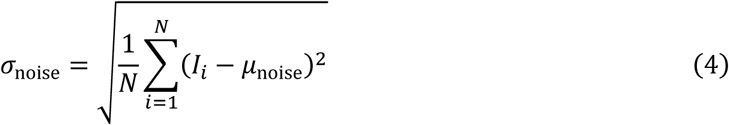

where I_i_ is the intensity of the i-th pixel and μ_noise_ is the mean intensity ‘off’ image.

SNR was defined as the ratio of the mean signal intensity (μ_sig_) in the noise standard deviation (σ_noise_):

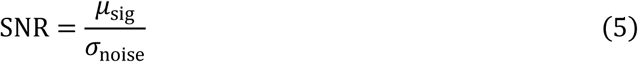

CNR was calculated to evaluate the visibility of details relative to the noise:

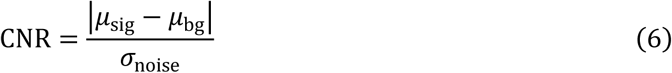

## Author contributions

J.Q. and T.Y. designed and built the optical system and the data acquisition software. J.Q. performed the data acquisition and analysis. F.G. provided support on biochemistry and live-cell sample preparation. M.A.P., J.Q. and T.Y. wrote the paper. M.A.P. supervised the whole study. All authors edited the paper.

## Data and materials availability

All data needed to evaluate the conclusions in this paper are present in the main text and/or the Supplementary Materials.

We thank S. Lee for assisting with editing of the paper.

---

**Supplementary Fig 1.**
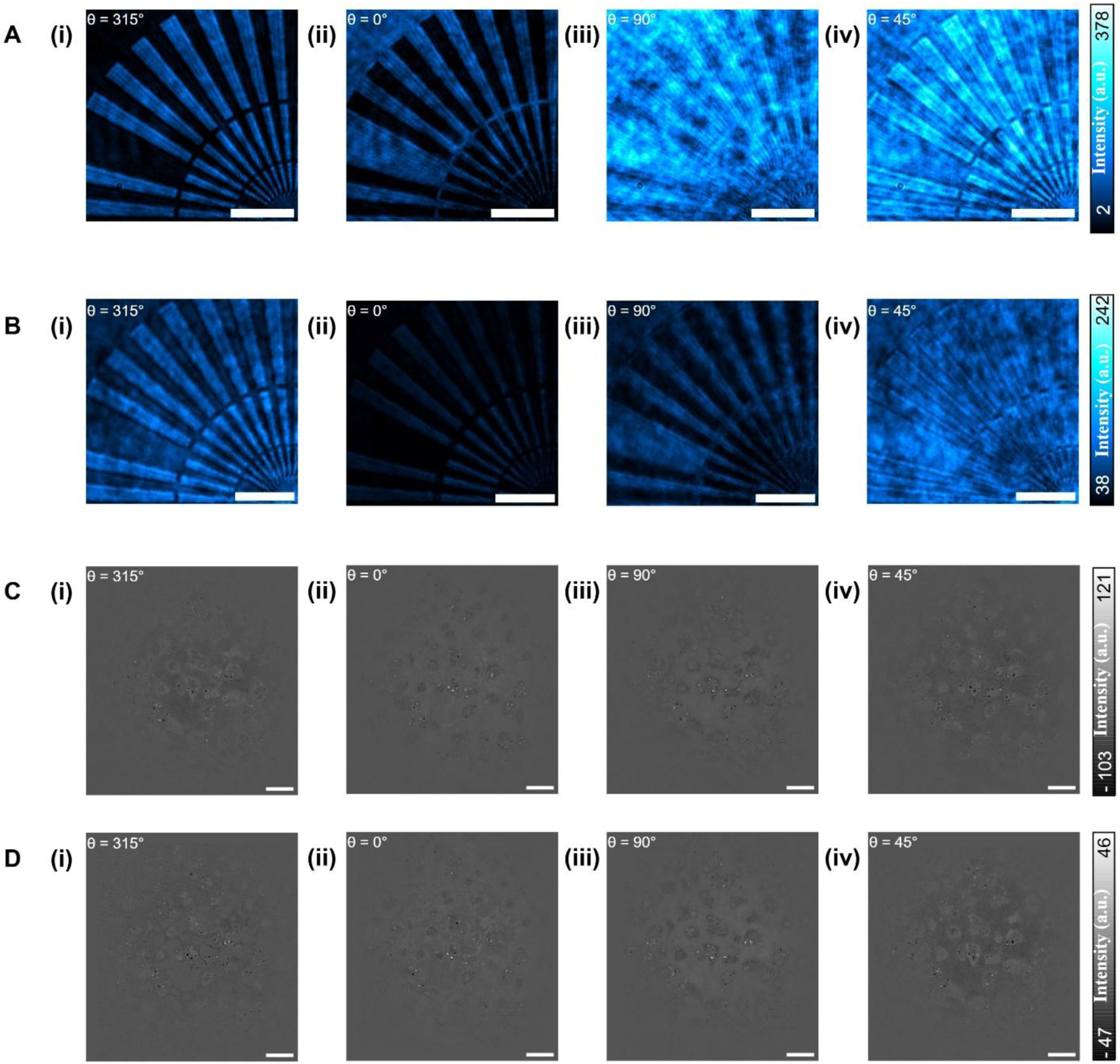
Comparison of phase-shifting module (PSM) images acquired by Instantaneous Phase-Shifting Optothermal Microscopy (IPSOM) and sequential Phase-Shifting Optothermal Microscopy (PSOM). **A.** PSM images of a focus star (nominal height 150 nm) acquired via IPSOM at polarization angles of 315° (i), 0° (ii), 45° (iii) and 90° (iv). **B.** PSM images of a focus star acquired via PSOM. **C.** PSM of fixed (day-6 differentiated) adipocytes acquired via IPSOM. **D.** PSM of fixed (day-6 differentiated) adipocytes acquired via PSOM. Scale bars: 50 µm.

**Supplementary Fig 2.**
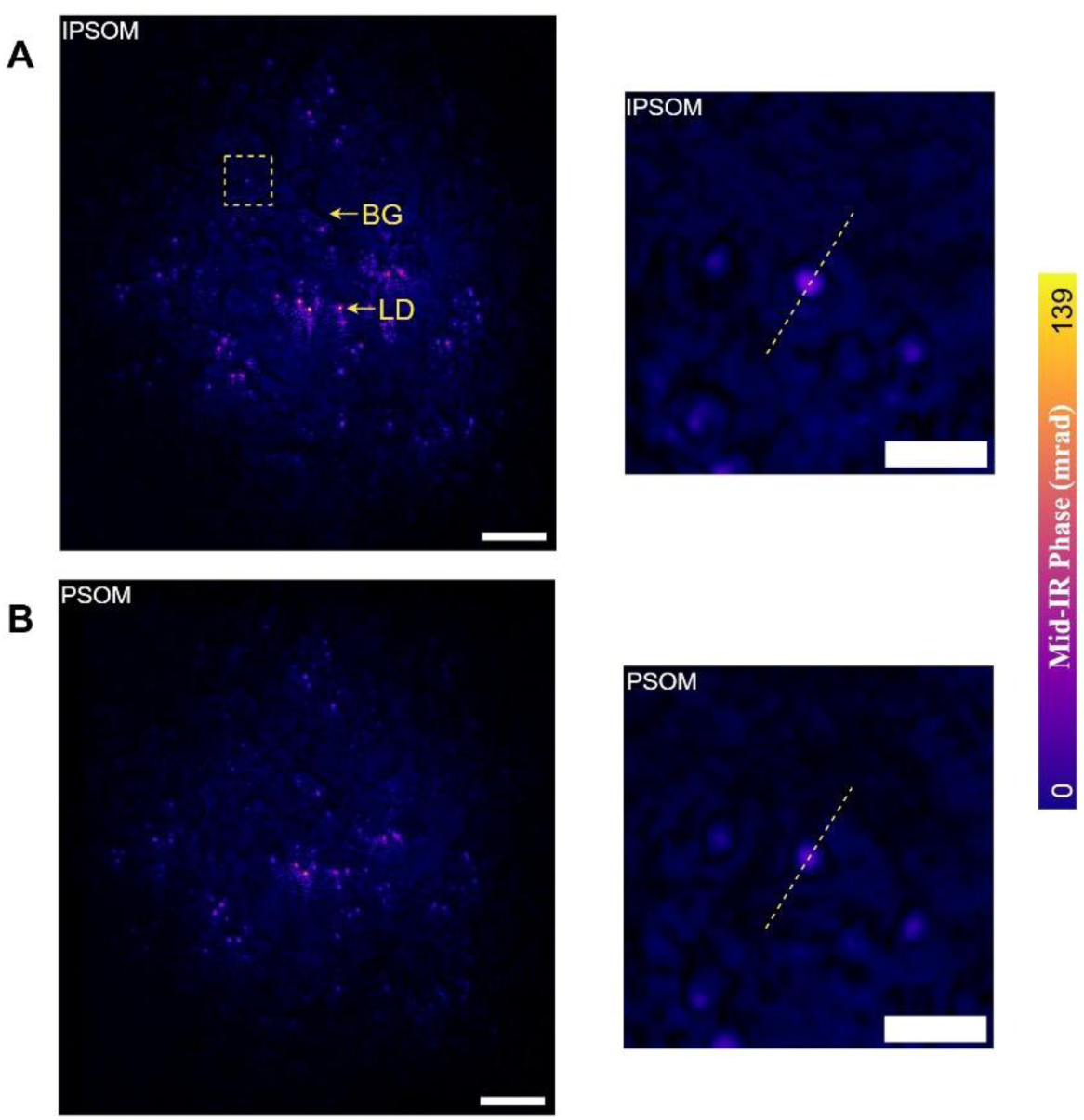
Comparison of micrographs of fixed differentiated adipocytes acquired using Instantaneous Phase-Shifting Optothermal Microscopy (IPSOM) and sequential Phase-Shifting Optothermal Microscopy (PSOM). **A.** IPSOM micrograph, showing a zoomed in version of the square area highlighting a small lipid droplet where the line profile is acquired from. **B.** PSOM micrograph and zoomed in highlight. Scale bars: 50 µm, 20 µm.

**Supplementary Fig 3.**
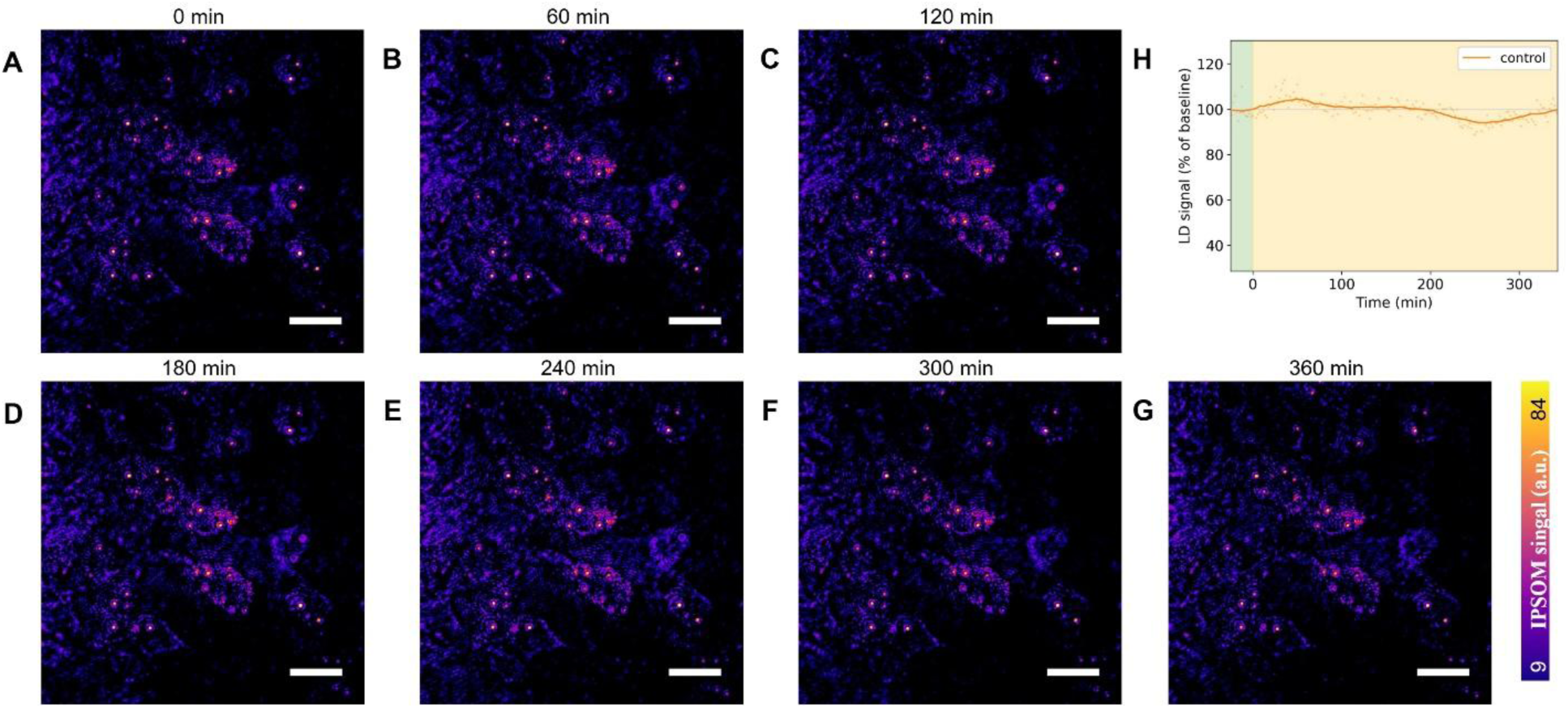
Monitoring lipid remodelling during lipolysis – control. **A-G.** Representative time series of Instantaneous Phase-Shifting Optothermal Microscopy (IPSOM) images of live differentiated (day 6) adipocytes acquired at 2850 cm^-1^. **H.** Lipid droplet (LD) signal, control signal stays relatively stable. Scale bars: 25 µm.

**Supplementary Fig 4.**
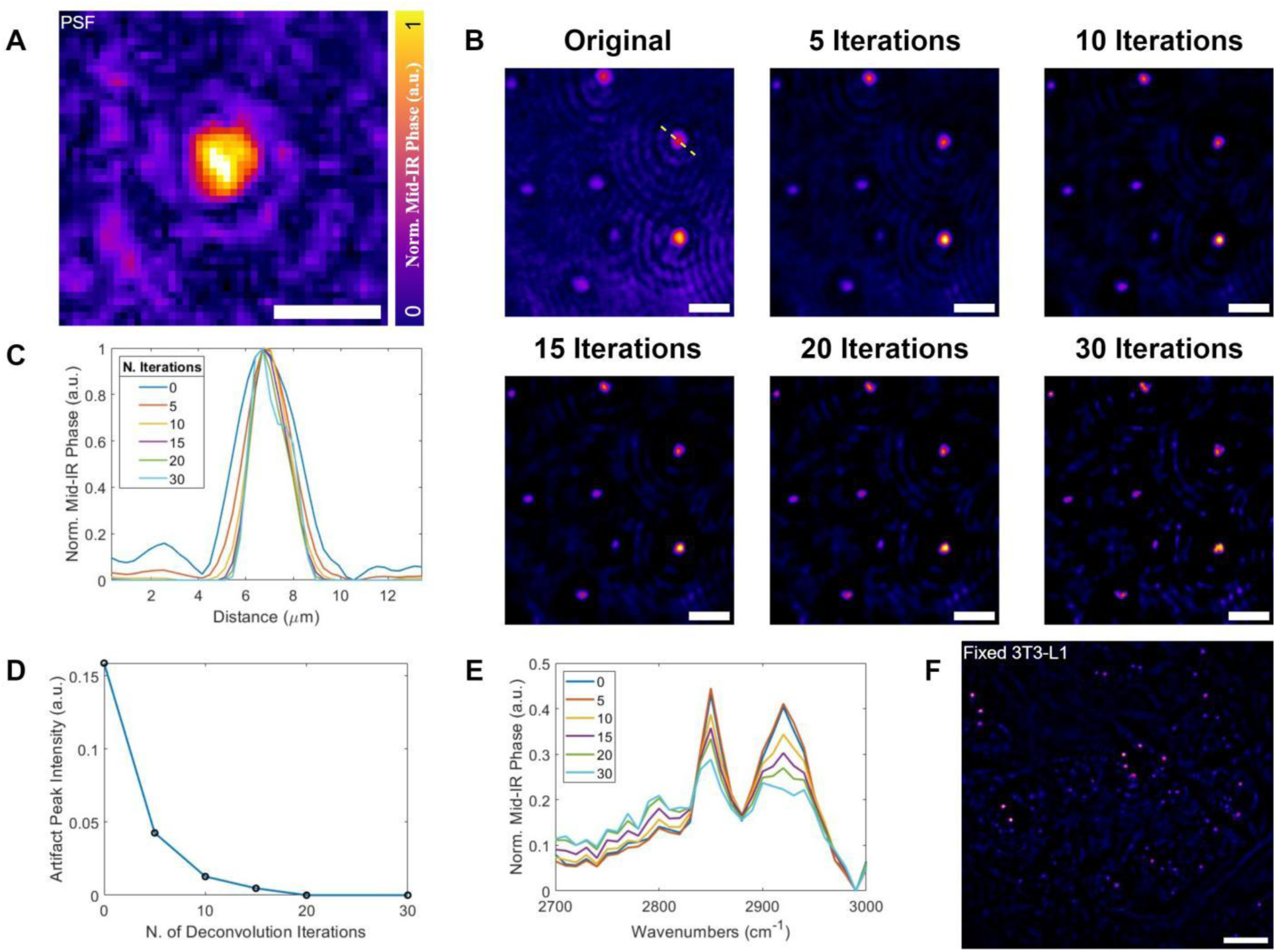
Results of deblurring by deconvolution using the mid-IR phase point spread function (PSF), triglyceride (TAG) droplets are imaged to extract the PSF. **A.** Mid-IR phase PSF. **B.** Deblurred images of TAG droplets, shows a comparison of the effect of increasing the number of deconvolution iterations. **C.** Line profile over the highlighted TAG droplet in **B**, artifacts are observed to be attenuated as the number of deconvolution iterations is increased. **D.** Plot of artifact peak intensity against the number of deconvolution iterations. **E.** Spectra of TAG after deblurring, the spectral shape changes after 10 or more deconvolution iterations are applied. **F.** Deblurring result of fixed differentiated (day 6) 3T3-L1 cells, the signal from lipid droplets are preserved while the signal from disk-like artifacts are attenuated. Scale bars: A: 5 µm, B: 10 µm, F: 25 µm.

